# *Ttc21b* is required in Bergmann glia for proper granule cell radial migration

**DOI:** 10.1101/181230

**Authors:** Ashley M. Driver, Christopher Shumrick, Rolf W. Stottmann

## Abstract

Proper cerebellar development is dependent on tightly regulated proliferation, migration,
and differentiation events. Disruptions in any of these leads to a range of cerebellar phenotypes from ataxia to childhood tumors. Animal models have shown proper regulation of *sonic hedgehog* (*Shh*) signaling is crucial for normal cerebellar architecture and increased signaling leads to cerebellar tumor formation. Primary cilia are known to be required for the proper regulation of multiple developmental signaling pathways, including *Shh*. *Tetratricopeptide Repeat Domain 21B* (*Ttc21b*) is required for proper primary cilia form and function and is primarily thought to restrict *Shh* signaling. Here we investigated a role for *Ttc21b* in cerebellar development. Surprisingly, *Ttc21b* ablation in Bergmann glia resulted in accumulation of ectopic granule cells in the lower/ posterior lobes of the cerebellum and a reduction in Shh signaling. *Ttc21b* ablation in just Purkinje cells resulted in a similar, phenotype seen in fewer cells, but across the entire extent of the cerebellum. These results suggest that *Ttc21b* expression is required for Bergmann glia structure and signaling in the developing cerebellum, and in some contexts, augments, rather than attenuates, *Shh* signaling.

## 1. Introduction

Development of the cerebellum includes significant postnatal growth and morphological change. The most numerous cell type in the mammalian brain is the cerebellar granule cell with estimates of approximately 100 billion cells in human (Andersen et al., 1992). In mouse and human, cerebellar granule cells are produced postnatally in the external granule layer (EGL). The post-mitotic granule cells migrate radially on Bergmann glial fibers to the interior of the cerebellum and ultimately form the inner granule layer (IGL). In humans, disrupted cerebellar development can result in a wide variety of structural malformations including cerebellar hypoplasia, failure of proper foliation, and tumor formation (Ten Donkelaar and Lammens, 2009).

*Sonic hedgehog* (*Shh*) signaling has become a well-recognized pathway involved in early cerebellar development. Granule cell proliferation and proper development of the Bergmann glial migratory tracts are both known to require sonic hedgehog (SHH) secreted from Purkinje cells (Dahmane and Ruiz i Altaba, 1999; Wallace, 1999; Wechsler-Reya and Scott, 1999; Xu et al., 2013). The downstream GLI transcription factors have dynamic expression patterns during early cerebellar development (Corrales et al., 2004).

Altered levels of *Shh* signaling have severe cerebellar developmental consequences and can cause tumor formation. Loss of *Shh* signaling results in a reduction of granule cell proliferation and cerebellar hypoplasia (Corrales et al., 2006; Lewis et al., 2004; Spassky et al., 2008). In addition to hypoplasia, conditional ablation of *Gli2* with an *En1-Cre* led to abnormal Purkinje cell and Bergmann glia morphology (Corrales et al., 2006). Deletion of *Gli1 and Gli2* leads to further disruptions in foliation and patterning (Corrales et al., 2006). In humans, impaired SHH signaling has been implicated in the cerebellar anomalies seen in both Joubert and Meckel Syndrome (Aguilar et al., 2012). Conversely, activation of SHH signaling via loss of *Patched*-1 or constitutively activated Smoothened mutations results in medulloblastoma formation (Goodrich and Scott, 1998; Hallahan et al., 2004; Han et al., 2009; Oliver et al., 2005; Yang et al., 2008).

The primary cilium is a non-motile sensory organelle that is now well established to be critical for proper modulation of the SHH pathway (Bangs and Anderson, 2017). Many proteins are trafficking through the primary cilium via intraflagellar transport (IFT). GLI proteins in particular require IFT through the primary cilium for proper processing into transcriptional repressor/activator forms (Liu et al., 2005; Tran et al., 2008). Loss of cilia and/or IFT leads to multiple defects affecting SHH signal transduction, including perturbation of GLI processing patterns. Consistent with this role in SHH signal transduction, loss of primary cilia causes cerebellar hypoplasia (Spassky et al., 2008) and modulates the effects of other mutations on tumor formation (Han et al., 2009).

Among the IFT proteins, *Tetratricopeptide Repeat Domain 21B* (*Ttc21b*; *Ift139*) is required for normal ciliary retrograde transport (Tran et al., 2008) and early forebrain patterning and development (Snedeker et al., 2017; Stottmann et al., 2009). Loss of *Ttc21b* has been shown to result in abnormal GLI3 processing, and thus increased Shh signaling (Stottmann et al., 2009; Tran et al., 2008). In humans, *TTC21B* is one of the most commonly mutated genes in ciliopathy patients, and variants are present in disorders of the cerebellum such as Joubert Syndrome (Davis et al., 2011). We hypothesized loss of *Ttc21b* during postnatal development may affect SHH signaling and cerebellar development. To investigate this and circumvent the perinatal lethality of *Ttc21b* mutants (Tran et al., 2008), we used the Cre-loxP system to separately ablate *Ttc21b* in Bergmann glia and Purkinje cells. Surprisingly, loss of *Ttc21b* resulted in ectopic granule cells in the posterior lobes of the cerebellum. Analysis of the affected lobes showed abnormal morphology of both Bergmann glia and Purkinje cells, as well as reduced SHH signaling. These results demonstrate a unique requirement for *Ttc21b* in cerebellar astrocytes and the first demonstration of reduced SHH signaling upon loss of *Ttc21b.*

## 2. Materials and Methods

### Mouse husbandry

All animals were maintained through a protocol approved by the Cincinnati Children’s Hospital Medical Center IACUC committee (IACUC2016-0098). Mice were housed in a vivarium with a 12-hour light cycle with food and water *ad libitum*. All mouse alleles used in this study were previously published: *Ttc21b*^*alien*^ is a null allele of *Ttc21b* (Tran et al., 2008), *Ttc21b*^*tm1a(KOMP)Wtsi-lacZ*^ (*Ttc21b*^*lacZ*^) for analysis of *Ttc21b* expression via the LacZ expression cassette (Tran et al., 2014), *Ttc21b*^*flox*^ (Snedeker et al., 2017; Tran et al., 2014); a Cre recombinase reporter allele B6.129(Cg)-*Gt*(*ROSA*)*26Sor*^*tm4(ACTB-tdTomato,-EGFP)Luo*^/*J* (*ROSA*^*dTom/EGFP*^) (Muzumdar et al., 2007); FVB-Tg(GFAP-cre)^25Mes/J^ (GFAP-Cre) (Zhuo et al., 2001) and B6.129-Tg(Pcp2-cre)^2M_pin_/J^ (Pcp2-Cre)(Barski et al., 2000). *Gli1*^*tm2AlJ*^/*J* (*Gli1*^*lacZ*^) mice were used to measure *Gli1* expression (Bai et al., 2002). For adult analysis, 4-month-old animals were sedated and transcardial perfusion with paraformaldehyde was performed using standard procedures. Brains were subsequently harvested and fixed for 48 hours in 4% paraformaldehyde at room temperature (RT) and embedded in paraffin. For postnatal brain collections, pups were sedated with isoflurane to minimize discomfort followed by decapitation. Brains were collected and fixed for 24 hours at 4°C and embedded in O.C.T. medium (Tissue Tek) for cryosectioning or in paraffin.

### Histology and Immunohistochemistry

For histological analysis, brain sections were collected on a Leica microtome at a thickness of 5μM. Sections were de-paraffinized, hematoxylin and eosin stained, and sealed using Cytoseal (Thermo Scientific). Images were obtained with a Zeiss Stereoscope (all paired images are shown at the same magnification). For DAB immunohistochemistry, sections were de-paraffinized and bleached with hydrogen peroxide. Sections were incubated overnight in primary antibody (NeuN, 1:500; Ki67 1:500) at RT. Sections were then washed in PBS and incubated for two hours in biotinylated swine-anti rabbit secondary antibody (1:500; Vector Laboratories) for 2 hours at RT. Sections were then washed in PBS and incubated in ABC solution (Vector Laboratories) for 1 hour at RT. Sections were then washed and visualized using DAB for 3 minutes followed by a PBS wash, ethanol dehydration, xylene incubation and mounted Cryoseal (Thermo). For fluorescent immunohistochemistry, sections were collected on a Leica cryostat at 12μM thickness and washed in PBS prior to antigen retrieval with sodium citrate buffer (as appropriate for each antibody). Sections were then washed with PBS and incubated in blocking buffer (PBS+ 4% normal goat serum and 0.1% Triton-X) for 1 hour at RT. Sections were incubated overnight at 4°C in primary antibody diluted in blocking buffer: Calbindin 1:1000 (Abcam, ab25085), GFAP 1:500 (Abcam, ab7260), pHH3 1:500 (SIGMA, H0412), and TTC21B 1:200(Novus Biologicals, NBP1-90416). Sections were then washed and incubated in Alexa Fluor-conjugated secondary antibodies diluted in blocking buffer (1:500, Life Technologies) followed by PBS washes and DAPI counterstain (1:1000) for 10 minutes. To visualize the GFP expressed by the *ROSA*^*dTom/EGFP*^ Cre reporter allele, sections were only incubated for 1 hour at RT with anti-GFP antibody (1:100, Life Technologies, A21311) followed by washes and DAPI counterstain. Sections were sealed with ProLong Gold (Life Technologies) and images were obtained on a Nikon C2 confocal microscope. All paired images were taken at the same magnification.

### Section in situ hybridization and lacZ staining

For RNA section *in situ* hybridization (SISH) and LacZ staining, sections were collected on a Leica cryostat at a thickness of 20μM. The in situ hybridization was done according to standard protocols. For LacZ staining sections were washed in PBS followed by fixation in cold 0.5% PFA on ice followed by PBS wash and incubation in LacZ buffer on ice. Sections were then incubated in X-Gal solution at 4°C for 1-3 days after which they were washed in PBS, counterstained in Eosin, and dehydrated. Sections were sealed with Cytoseal (Thermo) and images were taken on a Zeiss Stereoscope with all paired images taken at the same magnification.

## 3. Results

### 3.1 *Ttc21b is enriched in the Purkinje cell layer of the developing cerebellum*

We used a *Ttc21b*^*lacZ*^ allele to determine the pattern of *Ttc21b* expression in the developing cerebellum. LacZ staining from P1-P10 showed expression in the Purkinje cell layer (PCL,) and the the deep cerebellar nuclei (DCN: Fig 1. A-F). The Purkinje cell layer at these stages is comprised of both Purkinje cells and the cell bodies of Bergmann glia (Yamada and Watanabe, 2002). We performed immunohistochemistry to both verify the fidelity of the lacZ reporter expression, and to further define the cell type(s) expressing *Ttc21b*. At P1, we observed TTC21B in the cytoplasm of the Purkinje cells and in puncta consistent with primary cilia, a previously demonstrated site of *Ttc21b* expression (Fig 1. G-I). We also saw TTC21B in Bergmann glia (BG) cells highlighted by a GFAP antibody. In Bergman glia, the only TTC21B immunoreactivity was in puncta consistent with primary cilia, and not in the cytoplasm as was seen in the Purkinje cells. We confirmed this ciliary localization of TTC21B with three dimensional optical reconstructions (Fig 1. J-M). Thus, we conclude *Ttc21b* is expressed in both cells types in the early postnatal Purkinje cell layer, with cytoplasmic and ciliary localization in Purkinje cells, but only ciliary localization in Bergmann glial cells.

**Figure 1.**
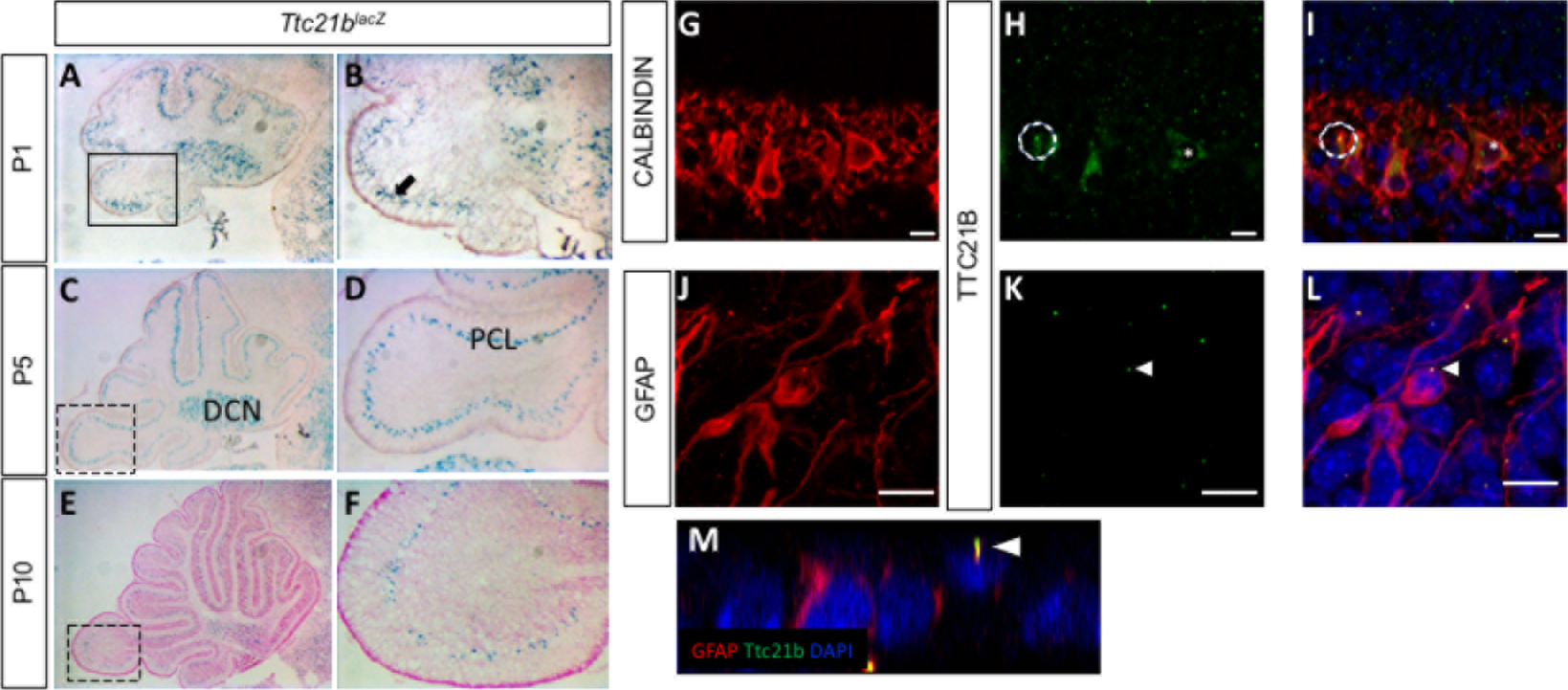
*Ttc21b* is expressed in the postnatal mouse cerebellum. (A-F) LacZ expression from the *Ttc21b*^*lacZ*^ reporter is evident in the Purkinje cell layer of the cerebellum from P1-P10. Immunohistochemistry for Calbindin in Purkinje cells (G) shows significant overlap with TTC21B (H,I). TTC21B is seen in both the cytoplasm (asterisk) and in cilia (e.g., circle) of Purkinje cells. TTC21B co-localization is also seen with the astrocyte marker GFAP (J) where TTC21B is found in primary cilia, but not the cytoplasm (K,L, arrowheads). Threedimensional reconstructions are consistent with a ciliary co-localization pattern (M, arrowhead). Boxes in A,C,E indicate areas of focus in B,D,F respectively. Scale bars in G-L represent 10 μm. n≥3 for LacZ staining, n≥2 for immunohistochemistry.

### 3.2. *GFAP-Cre ablation of Ttc21b results in ectopic granule cells in the cerebellum*

Given the known roles of SHH signaling in early postnatal cerebellar development and the role of *Ttc21b* in regulating Shh signaling, we sought to determine if *Ttc21b* was required in the developing cerebellum. To circumvent the perinatal lethality of *Ttc21b* null mutations (Tran et al., 2008), we used a conditional genetic approach and performed a genetic ablation using GFAP-Cre (Zhuo et al., 2001) (Fig S1. A-C). Consistent with previous reports, GFAP-Cre stimulated recombination was observed in the telencephalic hemispheres and in the cerebellum. To confirm the cell types exhibiting Cre activity, we used a Cre recombinase reporter allele, *ROSA*^*dTom/EGFP*^, in combination with immunohistochemistry. We see significant co-localization of GFP immunoreactivity indicating Cre activity with GLAST marking immature astrocytes (Fig S1. G-I), but no such co-localization in the Calbindin-positive Purkinje cells (Fig S1. D-F).

We genetically ablated *Ttc21b* from Bergmann glia by crossing *GFAP-Cre;Ttc21b*^*aln/wt*^ mice with *Ttc21b*^*flox/flox*^ animals to generate *GFAP-Cre;Ttc21b*^*flox/aln*^ mice lacking functional *Ttc21b* in the entire GFAP-expressing lineage. Histological analysis of *GFAP-Cre;Ttc21b*^*flox/aln*^ animals at P7 showed dense areas of granule cells in the nodular region of the cerebellum which were not seen in control animals (Fig 2. A,B, arrowhead; (White and Sillitoe, 2013). This phenotype was also observed in 4-month-old adult cerebella. Interestingly, the ectopic cells were found in the posterior cerebellum (lobes IX and X; Fig 2. C-F). We demonstrated these cells are differentiated granule cells with immunohistochemistry for NeuN as a marker for differentiated neurons. Ki67 immunostaining for proliferating cells indicated these are not mitotically active (Fig 2G-J). This suggests that the ectopic cells, while displaced, are not forming tumors as has been previously shown upon manipulation of SHH signaling in the cerebellum.

**Figure 2:**
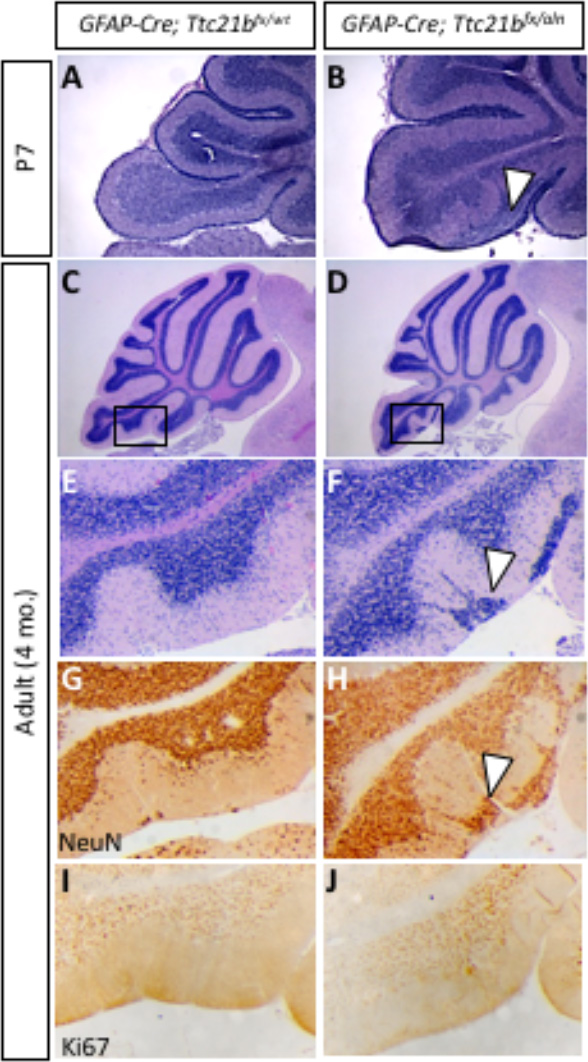
Ablation of *Ttc21b* from astrocytes results in ectopic granule cells in the cerebellum. H&E staining at P7 (A,B) and 4 months of age (C-F) show granule cells present in the outer layers of the cerebellum of *GFAP-Cre;Ttc21b*^*fx/aln*^ animals which are not found in controls (n>4). NeuN immunoreactivity indicates these ectopic cells are differentiated granule cells (G,H, arrowheads). Ki67 immunostaining for proliferative cells is negative (I,J) suggesting these are not proliferating tumors. n≥4 for adult histology, n<u>≥3</u> for adult DAB staining and P7 histology.

### 3.3 *Purkinje cell and Bergmann glial cell development are disrupted upon loss of Ttc21b*

We next sought to determine why the ectopic cells may be present in the *GFAP-Cre;Ttc21b*^*flox/aln*^ animals. We initially reasoned they may be granule cells that did not properly migrate from the site of formation in the external granule layer. This migration occurs along the radially oriented Bergmann glial fibers as part of normal granule cell development (Xu et al., 2013). To assess this hypothesis, we performed an immunohistochemical analysis of Bergmann glial cell development.

Immature Bergmann glia were specified in approximately the appropriate numbers and appear to have established radial fibers by P1 as shown by immunoreactivity for both Nestin and GLAST (Fig. 3A-D). Purkinje cells were specified and migrated to the appropriate position in the *GFAP-Cre;Ttc21b*^*flox/aln*^ cerebella at P1 (Fig 3. E,F). However, by P11 the *GFAP-Cre;Ttc21b*^*flox/aln*^ animals showed disrupted Bergmann glia fibers in the lower lobes of the cerebellum failing to form appropriate scaffolds for migration (Fig 3. G,H). Interestingly, the anterior lobes where we did not note ectopic cells appeared to have normal Bergmann glia in the *GFAP-Cre;Ttc21b*^*flox/aln*^ animals (Fig 3. K,L). At P11, the Purkinje cell bodies were irregularly aligned and the dendritic arbors were more extensive in the *GFAP-Cre;Ttc21b*^*flox/aln*^ animals as compared to control (Fig 3. K,L).

**Figure 3:**
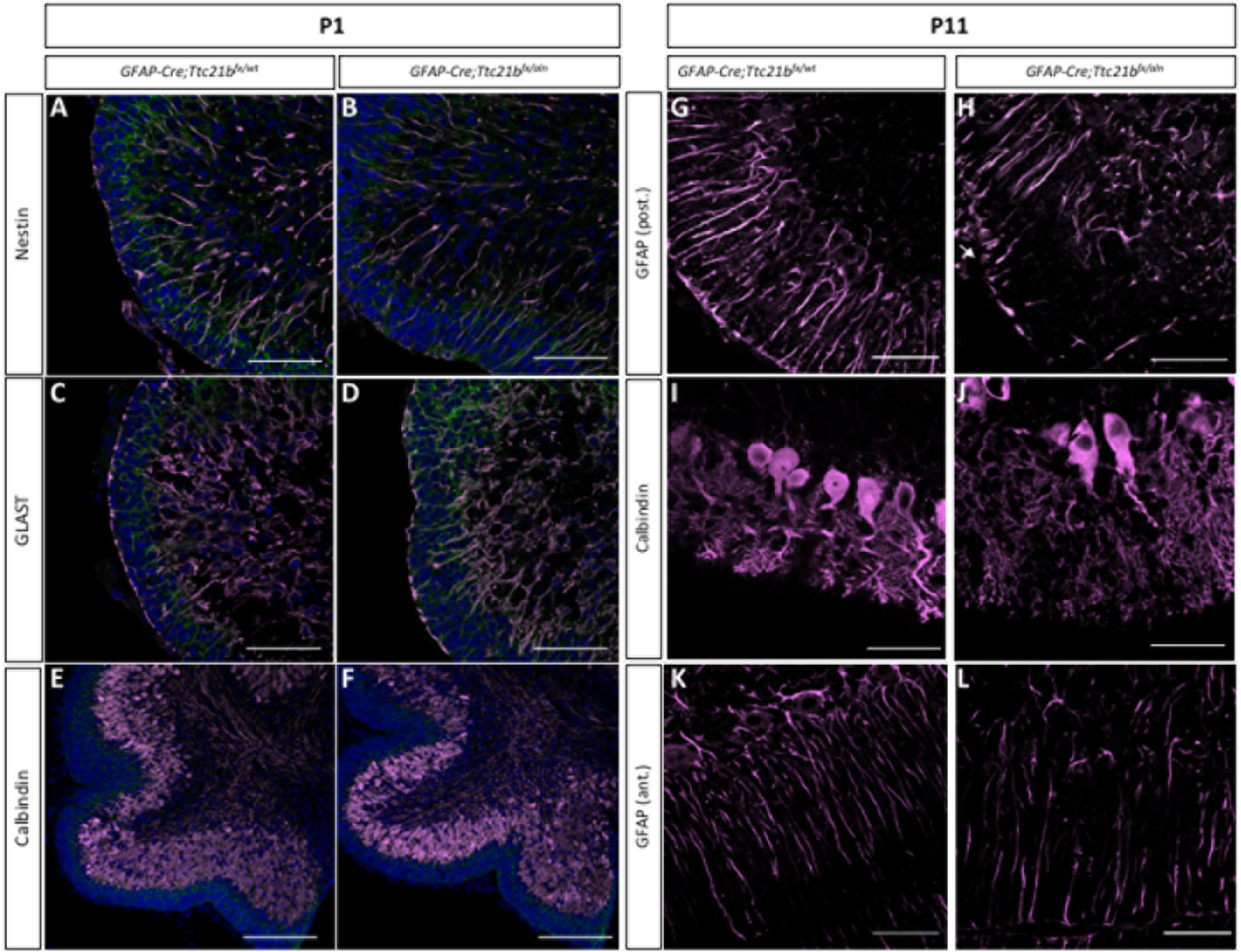
Purkinje cells and Bergmann glia development is disrupted in the *GFAP-Cre;Ttc21b*^*fx/aln*^ cerebella. (A-D)Immunostaining for immature astrocytes at P1 with Nestin (A,B) and GLAST (C,D) appears similar between control and *GFAP-Cre;Ttc21b*^*fx/aln*^ animals. (E,F) Calbindin immunostaining highlights the presence of Purkinje cells at P1 in both control and *GFAP-Cre;Ttc21b*^*fx/aln*^ animals. (G-J) P11 GFAP-positive mature astrocytes (G,H) and Calbindin-positive Purkinje cells (I-L) appear abnormal in the lower lobes of the *GFAP-Cre;Ttc21b*^*fx/aln*^ cerebella. (K,L) Anterior lobes still show normal Bergmann glia morphology in all animals. n= 3 for each genotype. Scale bars 50μm.

### 3.4 *SHH signaling is reduced in GFAP-Cre;Ttc21b^*flox/aln*^ cerebella*

As the primary cilium is a major hub for Shh signaling and loss of *Ttc21b* has been shown to affect SHH signaling, we then investigated SHH pathway activity in the *GFAP-Cre;Ttc21b*^*flox/aln*^ animals with the *Gli1*^*lacZ*^ allele (Lee et al., 1997). *Gli1*^*lacZ*^ expression was reduced in the Purkinje cell layer of *Gli1*^*lacZ*^;*GFAP-Cre;Ttc21b*^*flox/aln*^ animals (Fig 4. A-D, arrow). Since *Gli1* is not expressed by Purkinje cells, these results suggest a loss of SHH signaling activity in Bergmann glia (Corrales et al., 2004). RNA section *in situ* hybridization for another *Shh* transcriptional target, *Patched-1 (Ptch1)* also showed reduced expression in the Purkinje cell layer of the *GFAP-Cre;Ttc21b*^*flox/alien*^ animals as compared to controls at P11 (Fig 4. E,F,arrowhead). Expression of the *Shh* itself, however, was at similar levels between *GFAP-Cre;Ttc21b*^*flox/aln*^ and control animals, suggesting that Purkinje cell expression of the secreted SHH ligand is not affected (Fig 5. G,H). Taken together, these data show that *Shh* signaling is reduced in the adult *GFAP-Cre;Ttc21b*^*flox/aln*^ cerebellum, a striking difference to previous observations of *Ttc21b* function in the nervous system.

**Figure 4:**
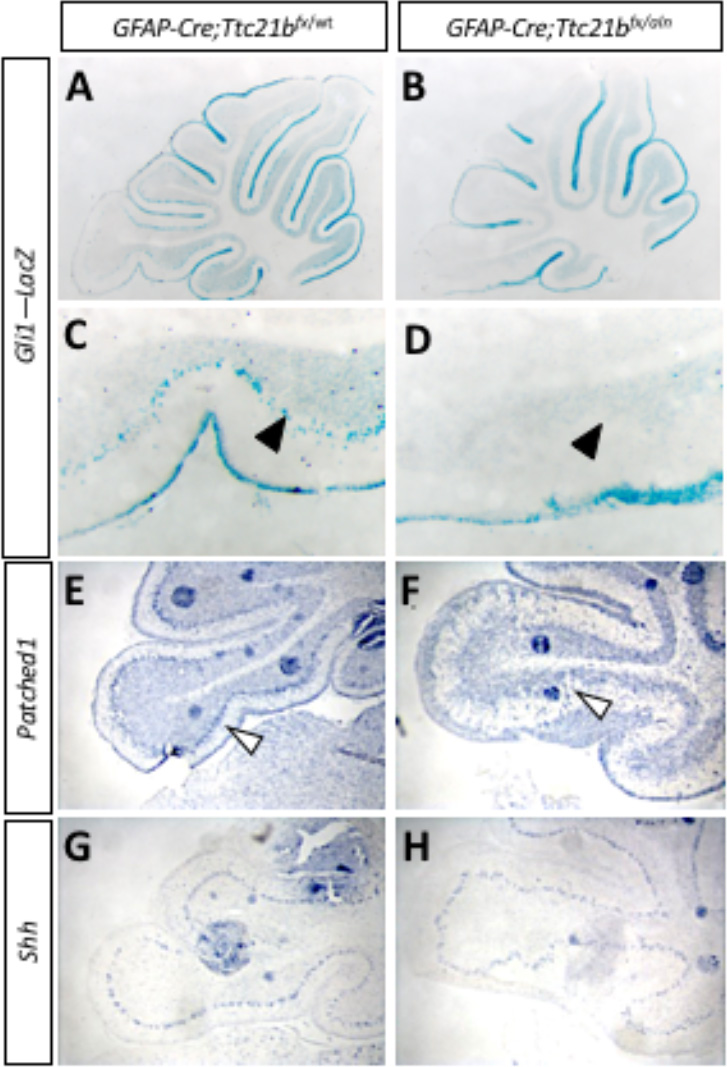
Shh signaling is reduced in *GFAP-Cre;Ttc21b*^*fx/aln*^ cerebella. (A-D) P11 *Gli1*-LacZ activity is reduced in in the Purkinje cell layer (arrows) of the *GFAP-Cre;Ttc21b*^*fx/aln*^ animals (B,D) as compared to control (A,C). (E,F) *Patched1* expression is also reduced in the *GFAP-Cre;Ttc21b*^*fx/aln*^ cerebella at P11 (F) compared to control (E), especially in the Purkinje cell layer (open arrowhead). (G,H) *Shh* expression appears similar between control (G) and *GFAP-Cre;Ttc21b*^*fx/aln*^ (H) but the organization of Shh cells again appears disorganized in the lower lobe of the cerebellum in ablated animals.

**Figure 5:**
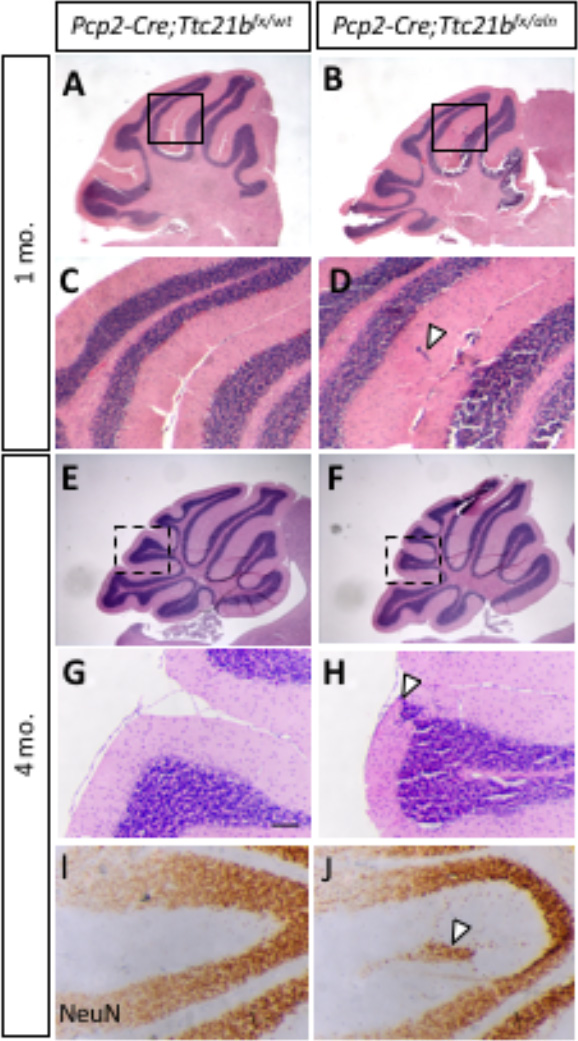
Ablation of *Ttc21b* with *Pcp2-Cre* results in similar, but milder, ectopic granule cell phenotype. (A,B) Small areas of displaced granule cells (arrowhead) are present in *Pcp2-Cre;Ttc21b*^*fx/aln*^ brains at 1 month of age. (B,D: boxes in A,B show areas highlighted in C, D). These cells are still visible at 4 months of age in the *Pcp2-Cre;Ttc21b*^*fx/aln*^ animals (H, arrowhead) and are immuno-positive for NeuN indicating these cells are differentiated granule cells (J, arrowhead). n≥3 for each genotype.

### 3.5 *Purkinje cell ablation of Ttc21b results in milder ectopic granule cell phenotype*

Since we observed *Ttc21b* expression in both Purkinje cells and Bergmann glia, we performed a similar genetic ablation of *Ttc21b* in the Purkinje cells with the *Pcp2-Cre* (Barski et al., 2000) (Fig. S2). At 1 month of age, histological analysis again revealed small areas of ectopic cells in the outer layer of the *Pcp2-Cre;Ttc21b*^*flox/aln*^ cerebella (Fig 5. A-D, arrowhead). This was also periodically seen in 4-month-old *Pcp2-Cre;Ttc21b*^*flox/aln*^ as well (Fig 5. E-H, n=4). Immunohistochemistry for NeuN again suggesting these cells were differentiated granule cells which failed to migrate to the IGL (Fig 5. I,J). Interestingly, this phenotype was less penetrant that in *GFAP-Cre;Ttc21b*^*flox/aln*^ animals, but was not solely restricted to the posterior lobes.

## 4. Discussion

Here we have demonstrated a role for *Ttc21b* in the both the Bergmann glia astrocytes and Purkinje cells of the developing mouse cerebellum. Using a *GFAP-Cre* genetic ablation of *Ttc21b* we showed a few granule cells are retained in the outer layer of cerebellum, specifically in lobes IX and X. This appears to be a failure of granule cell migration as the Bergmann glia, substrates for tracts for granule cell migratory, have reduced radial projections in the lower lobes of the ablated animals. In contrast, a *Pcp2-Cre* Purkinje cell specific ablation has a similar, but much less severe, phenotype, with only sparse areas of ectopic granule cells. Additionally, while GFAP-Cre recombination occurs in Bergmann glia, we noted effects on Purkinje cell development, supporting previous suggestions that a critical interaction between these two cells types is required for proper Purkinje cell dendritic development. We also observed ablation of *Ttc21b* resulted in a reduction in SHH signaling in the Purkinje cell layer, which is the first time we have a reduction on the Shh pathway upon loss of *Ttc21b*.

The radial fibers of the Bergmann glia serve as primary migratory tracts for granule cells in the developing cerebellum (Xu et al., 2013) and disruptions of these radial fibers have been previously shown to be associated with ectopic granule cells (Frick et al., 2012; Inouye et al., 1992). We only observed disruption of mature Bergmann glia in the affected lobes of the mutant suggesting this is a primary cause of the failed granule cell migration. Additionally, we noted abnormalities in Purkinje cell morphology in *GFAP-Cre;Ttc21b*^*flox/aln*^ animals consistent with numerous interactions between Bergmann glia and Purkinje neurons (Bellamy, 2006; Lordkipanidze and Dunaevsky, 2005; Wang et al., 2012). The ectopic granule cell phenotype was much less severe in the *Pcp2-Cre; Ttc21b*^*flox/aln*^ animals. This could suggest differing, cell-specific requirements for *Ttc21b*. Indeed, patients with *TTC21B* variants have shown cytoskeletal abnormalities in human podocytes in addition to abnormal cilia suggesting multiple roles for *TTC21B* (Huynh Cong et al., 2014). Additionally, *Pcp2-Cre* recombination does occur later in development than GFAP-Cre (P5) and may not completely ablate *Ttc21b* until after many cells have already completed migration.

Prior reports of *Ttc21b*^*aln/aln*^ mutants have shown increased SHH signaling in addition to abnormal GLI processing. Embryonically, loss of *Ttc21b* led to increased translation and abnormal proteolytic processing of GLI3 (Tran et al., 2008). Inappropriate SHH activity has also been demonstrated in embryonic forebrain of *Ttc21b*^*aln/aln*^ mutants with abnormal GLI3 processing (Stottmann et al., 2009; Tran et al., 2008). In a postnatal deletion of *Ttc21b*, increased GLI2 activity led to polycystic kidneys (Tran et al., 2014). In contrast to these findings we report a reduction in downstream *Patched* and *Gli1*, consistent with reduced Shh signaling.

While *GFAP-Cre* should lead to loss of *Ttc21b* throughout the cerebellum, we only observed the ectopic granule cell phenotype in the posterior lobes. GLI2 is enriched in the Purkinje cell layer in the posterior lobes of the developing cerebellum around P5 and has been shown to be the major effector of positive SHH signaling (Corrales et al., 2004). While all three GLI transcription factors are expressed in the developing cerebellum, *Gli1* becomes restricted to the Bergmann glia in the Purkinje cell layer (Corrales et al., 2004). However, it is unlikely that the phenotype reported here is solely due to an effect on *Gli1* as *Gli1* homozygous null mutants have a phenotypically normal cerebellum (Park et al., 2000). Both *Gli2* and *Gli3* null mice have abnormal cerebellar development, with primary defects in foliation (Blaess et al., 2008). Additionally, loss of *Gli2* has been shown to result in abnormal Bergmann glia morphology and Purkinje cell alignment (Corrales et al., 2006). Primary cilia are required for the proper processing of GLI2 protein into a transcriptional activator and GLI3 protein into a transcriptional repressor form. The loss of normal cilia function may have different effects on SHH signaling in different tissues depending on which GLI protein has a more prominent role at that particular time in development. Given the enriched expression of *Gli2* in the same area as the ectopic cells and its role in stimulating SHH signaling, the reduced SHH noted here in the postnatal cerebellum may be more reflective of the loss of the GLI2 activator form than the GLI3 repressor.

The role of both ciliary genes and SHH signaling is a rapidly expanding field of study in human cerebellar development. SHH signaling is dysregulated in cells from Joubert syndrome and Meckel syndrome patients (Aguilar et al., 2012). Interestingly, *TTC21B* variants are associated with Joubert syndrome patients (Romani et al., 2013). Our study shows *Ttc21b* has a role in the glial cerebellar cells and loss of *Ttc21b* in different tissues can lead to increased and decreased SHH signaling.

## Supplementary Materials

The following are available online at www.mdpi.com/link, Figure S1: GFAP-Cre recombination occurs in the Bergmann glia of the cerebellum, Figure S2: Pcp2-Cre recombination occurs in the Purkinje cells of the cerebellum.

## Acknowledgments

This work is supported by the National Institutes of Health (R01 NS085023).

### Author Contributions

A.D. and R.W.S. conceived and designed the experiments; A.D. and C.S. performed the experiments; A.D., C.S, and R.W.S. analyzed the data; A.D. and R.W.S. wrote the paper

### Conflicts of Interest

The authors declare no conflict of interest. The funding sponsors had no role in the design of the study; in the collection, analyses, or interpretation of data; in the writing of the manuscript, and in the decision to publish the results.

